# Simulation and reconstruction ofmetabolite-metabolite association networks usinga metabolic dynamic model and correlation based-algorithms

**DOI:** 10.1101/460519

**Authors:** Sanjeevan Jahagirdar, Maria Suarez-Diez, Edoardo Saccenti

**Affiliations:** Laboratory of Systems and Synthetic Biology, Wageningen University & Research,Stippeneng 4, 6708WE Wageningen, the Netherlands

## Abstract

Biological networks play a paramount role in our understanding of complex biological phenomena and metabolite-metabolite association networks are now commonly used in metabolomics applications. In this study we evaluate the performance of several network inference algorithms (PCLRC, MRNET, GENIE3, TIGRESS and modifications of the MRNET algorithm, together with standard Pearson’s and Spearman’s correlation) using as a test case data generated using a dynamic metabolic model describing the metabolism of arachidonic acid (consisting of 83 metabolites and 131 reactions) and simulation individual metabolic profiles of 550 subjects. The quality of the reconstructed metabolite-metabolite association networks was assessed against the original metabolic network taking into account different degrees of association among the metabolites and different sample size and noise levels. We found that inference algorithms based on resampling and bootstrapping to perform better when correlations are used as indexes to measure the strength of metabolite-metabolite associations. We also advocate for the use of data generated using dynamic models to test the performance of algorithms for network inference since they produce correlation patterns which are more similar to those observed in real metabolomics data.

## INTRODUCTION

Biological networks and network analysis are rapidly becoming fundamental tools for the investigation and the interpretation of metabolomics data within a systems biology context.^1^ A biological network can be defined as a system of interconnected biological sub-units or nodes: in metabolomics the nodes are metabolites measured in biological samples, and their association is defined on the base of their similarity, which is characterized by a similarity measure.^2^ Correlation is usually used as an index to measure the similarity of metabolite profiles and biological information can be derived from both the sign and the magnitude of the correlation: strong positive correlation can indicate an equilibrium condition or enzyme dominance, while strong negative correlation can indicate the presence of a conserved moiety.^3^

Metabolite-metabolite association networks can be used to describe comprehensibly a given status of a biological system or to compare the systems across multiple conditions; in fact, the metabolites association patterns can change upon perturbation of the system, induced under experimental control, or because of the onset of pathological conditions, under the assumption that differences and commonalities in the biological processes are reflected in the characteristics of the reconstructed networks.^4^

In recent times, thanks to the knowledge flow between the metabolomics and the systems biology communities, methods originally developed for the inference of gene regulatory networks have been proposed, extended and adapted to investigate the patterns of association among metabolites.^5–9^

In a previous study^10^ we briefly compared the performance of these algorithms on data with known correlation structures and generated using standard multivariate approaches available within computational environments like Matlab^11^ and R.^12^ However, we noted that generating synthetic data representing the patterns of associations between metabolites and their distributions as observed in real biofluid samples is a non trivial problem. Consider for instance blood that collects molecules from all different tissues in the body: a modeling framework to simulate concentrations of blood metabolites would need to account for reactions and metabolic pathways together with their interconnections and for the uptake and release in and from different organs, tissues, and cell types, forming a system governed by highly non linear dynamics.

For these reasons, generating data to test network inference algorithms using standard multivariate approaches, although straightforward, may not be the most appropriate strategy. In the present study we used a (simplified) model of metabolism to generate metabolite concentration profiles with physiologically plausible distributional and correlation (association) patterns.

This metabolic model is a network structure of interdependent variables (*i.e.* metabolites) that can be described using a system of ordinary differential equations (ODE’s). In this way the time course of metabolite concentrations are described as a function of rate laws that account for both enzymatic and non enzymatic reactions, implementing Michaelis-Menten^13^ and mass action^14^ kinetics. As a test case we reconstructed the metabolic pathway for the degradation of arachidonic acid (AA) making use of previous knowledge available in literature and biological databases. We used this dynamic model, consisting of 83 metabolites and 131 reactions, to generate physiologically plausible metabolite concentration profiles simulating data for 550 individuals, introducing inter-individual variability by randomly varying the kinetic constants of the model.

We used this data to test the performance of several algorithms to reconstruct metabolite-metabolite association networks, such as: PCLRC (Probabilistic Context Likelihood of Relatedness on Correlation),^7^ MRNET (Multicast REduction NETwork),^15,16^ GENIE3 (Gene Network Inference with Ensemble trees),^17^ TIGRESS (TRustfull Inference of Gene REgulation using Stability Selection)^18^ and modification of the MRNET algorithm, together with standard Pearson’s and Spearman’s correlation, by comparing the reconstructed networks with the structure of the original AA metabolic model.

Our results show the validity of such computational approach and indicates that methods based on resampling or non linear decision tress like Random Forest are be better suited for the reconstruction of metabolite-metabolite association networks than methods based on linear regression. Moreover, our approach also allows to evaluate to what extent high (partial) correlation coefficients correspond to known metabolic reactions, as it has also been shown,^3,5^ and we will also show, that metabolites directly linked in a reaction step may not exhibit any correlation.

## MATERIALS AND METHODS

### Dynamic model of Arachidonic Acid metabolism

Arachidonic acid (AA) and its derivatives (eicosanoids) have been extensively studied in the last 30 years due to their activity as signaling molecules and their role in the cross-regulation of the inflammatory response in the human body.^19–21^

Here we have developed a dynamic model of AA degradation by retrieving information about metabolites and eiconasoids involved in AA metabolism and associated reactions from the following databases (see Figure 1 for an overview and the Supporting material (Tables S1 and S2) for a complete list of metabolites): Recon2.2,^22^ the Human Metabolome Datababase (HMDB),^23–26^ the Kyoto Encyclopedia of Genes and Genomes (KEGG)^27–29^ and the Chemical Entities of Biological Interest database (ChEBI).^30–33^ The data retrieved from these databases included metabolite names, identifiers and IUPAC names, metabolite concentrations in normal physiological conditions in humans, reaction descriptions (substrates, products, cofactors and stoichiometry) and list of associated enzymes. Kinetic data, such as *K_m_* values for enzymes and *k_i_* values as reaction/equilibrium constants, were collected from BRENDA^34^ and SABIO-RK.^35^

**Figure 1.**
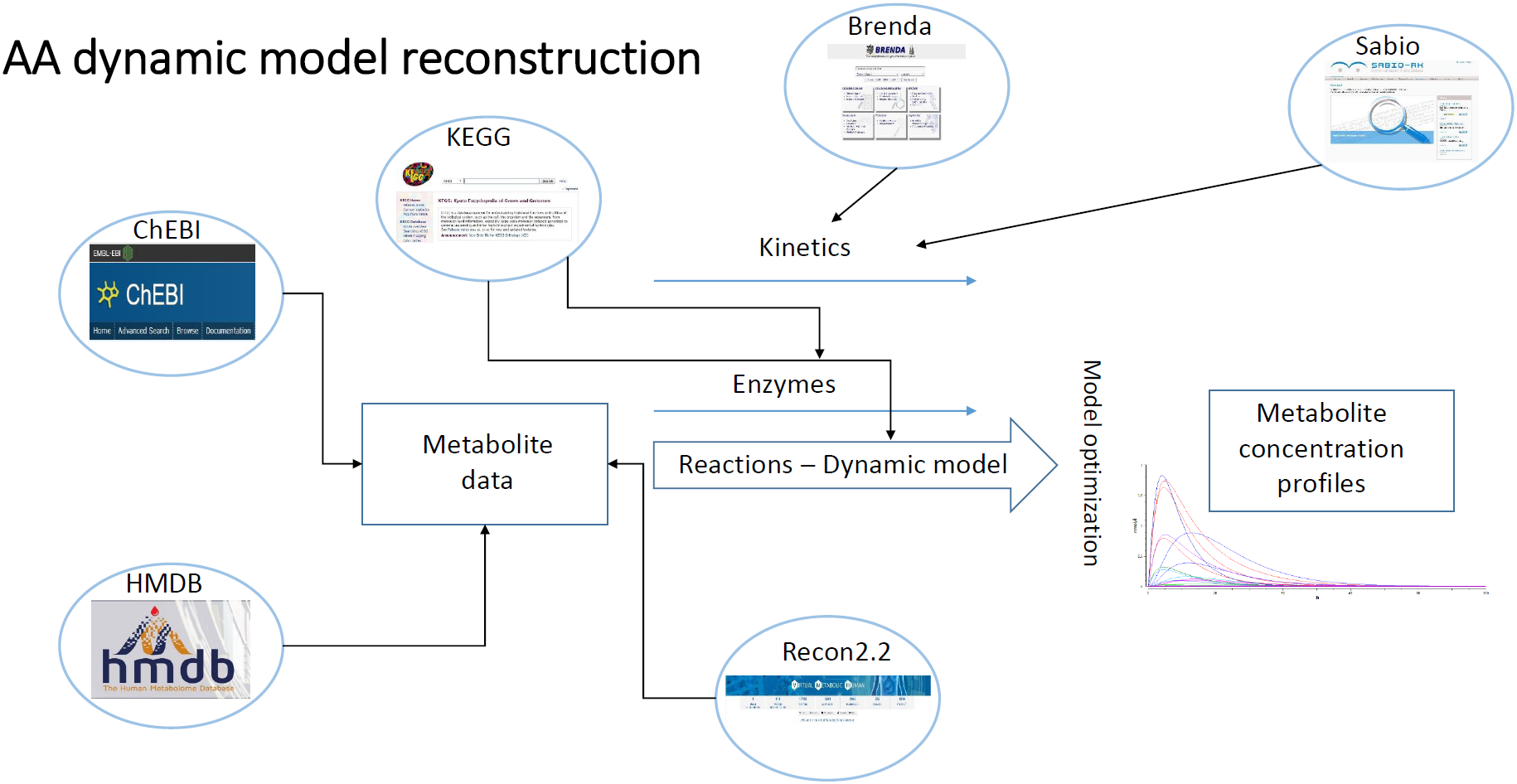
Overview of the strategy for the reconstruction of the the dynamic model for arachidonic acid (AA) metabolism. The databases used are indicated together with the type of information retrieved.

The reconstructed AA metabolic pathway is depicted in Figure 2. In this model, the starting point is activation of phosphatidylcholine (PC) that leads to AA synthesis and formation of AA-derived eicosanids such as prostaglandins, leukotrienes, thromboxanes and prostacyclines.^36,37^ The model accounts for concentrations of 83 metabolites and includes 131 reactions. An ODE was written for each metabolite in the model to account for synthesis, conversion and (when needed) degradation. Enzymatic reactions were described using Michaelis-Menten kinetics:^13^

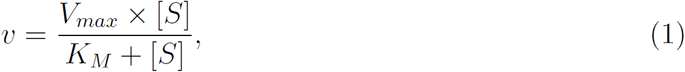

**Figure 2.**
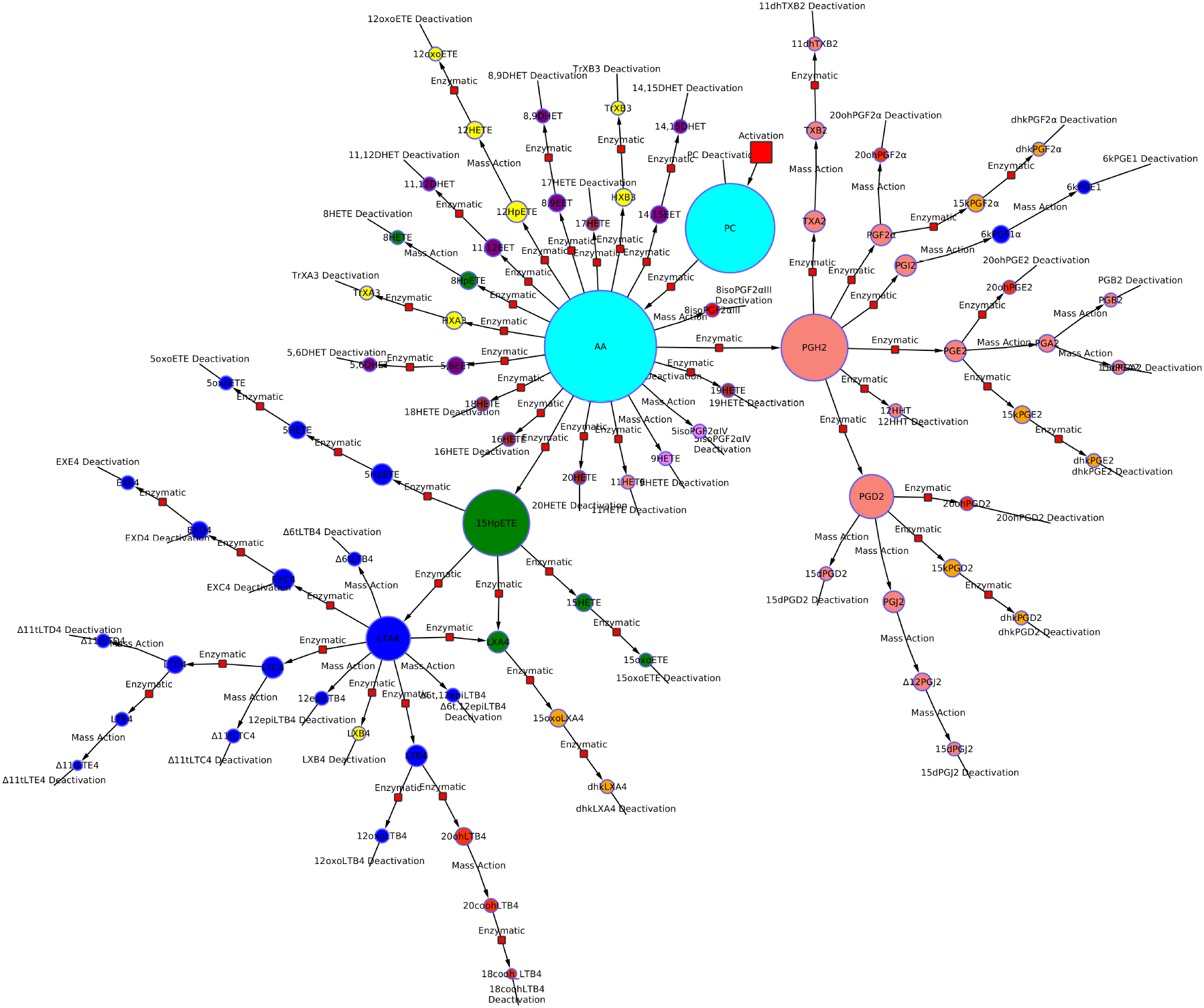
Graphical representation of the AA metabolic pathway. The enzymatic or nonenzymatic (mass action) nature of the reaction kinetic is indicated. The model comprises 131 reactions and 83 metabolites. The ODEs governing the dynamic model, metabolite desscription and reaction details are given in the supplementary material.

where υ is the rate of reaction, *V_max_* is the maximum reaction rate achieved by the system, [*S*] is substrate concentration, and *K_M_* is the Michaelis-Menten constant. Non-enzymatic reactions were described by simple mass action laws, where the rate of the reaction, υ is proportional to substrate concentration, [*S*], (*υ* = *k_i_* × [*S*].) and with the proportionality constant, *k_i_*, being the equilibrium constant of the reaction.

It should be noted that we have greatly simplified the reactions in this pathway and we have not explicitly described co-factors such as ATP or NADP, instead their impact is implicitly incorporated in the values of unknown kinetic constants through the fitting to measured data (see below).

A full description of the model (and ODEs) together with the description and values (when available) of all kinetic constants^25,38–75^ are given in the Supplementary Material S1 (Tables S1 and S2); the Supporting material S2 contains the metabolic model in SBML format.

As an example we provide here description of formation and degradation of AA, with the conversion of PC into AA: this is modelled as an ODE coupling the concentrations of each metabolite ([AA] and [PC] respectively):

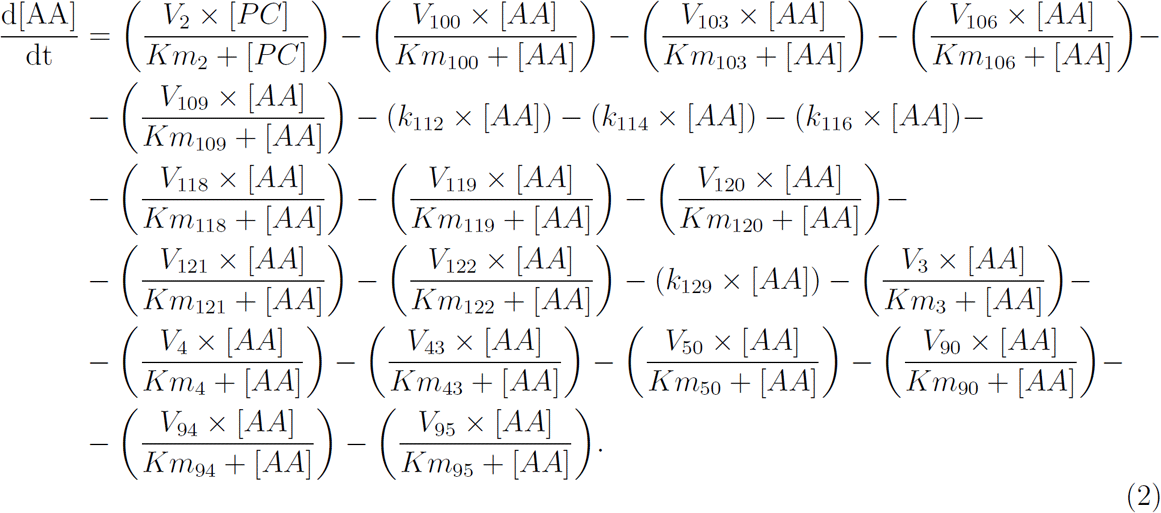

The model was constructed in a single compartment with a 1L volume so that concentration changes in metabolites (in nmol/L) due to small volume changes could be considered negligible.

### Optimization of parameters in the model

Not all reaction parameters for the reconstructed AA metabolic model (*k_i_*, *K_m_* and reaction rates) were available in literature. Missing parameters where estimated by identify sets of parameters maximizing the agreement between simulated concentrations (at steady state) and metabolite concentrations derived from literature (see Table S1 and S2 in the Supporting material S1 for a list of steady state concentrations and associated references). The optimization was done using the Genetic algorithm approach implemented in the simulation software COPASI^76^ using 2000 generations and each time choosing the 20 best performing sets of parameters.

### Simulation of individual metabolite concentration profiles

The metabolic model of AA metabolism was used to generate physiologically plausible metabolite concentration profiles for a pool of *N* individuals. Initial concentration of all metabolites was set to 0 except for phosphatidylcholine. Inter-individual variability (i.e. different profiles) was simulated by varying the *Km* and the *k_i_* constants and the reaction rates υ for all reactions in the model. Values for *Km*, *k_i_* and *υ* were sampled from an uniform distribution with lower and upper bounds set to the reference values ±10%. For instance, the *j*-th individual, the values of *k_i_*, *Km* and *υ* for a particular reaction were defined as

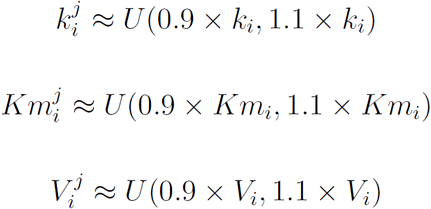

The reference values used for *k_i_*, *Km_i_* and *V_i_*, together with the associated references are given in Table S1 in the Supporting material.

Long term simulations showed the system (with optimal parameters) to achieve a steady state for all metabolites at approximately 70 hours after activation of the AA conversion. This time is consistent with previously reported experimental data.^77, 78^ For each of the *N* individuals, a time course of 100h was simulated and data was collected at two time points, 10 and 90 hours after induction of AA conversions. These correspond to non-steady state (NS) and steady state (SS) conditions.

### Noise addition

Gaussian uncorrelated noise with zero mean and variance-covariance matrix **Σ** = **I** (with **I** a 83 × 83 identity matrix was added to data matrices **X**_1_ and **X**_2_ to simulated experimental noise:

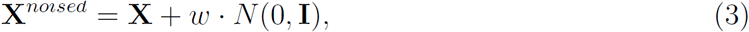

where the noise intensity levels *w* correspond to signal-to-noise ratios (SNR) of 1%, 5%, 10% and 15%. SNR is defined as the ratio of median peak intensity to the intensity of additive noise. X represents the data matrices containing the metabolite concentration profiles.

### Data overview

Overall, we generated *N* = 550 metabolites profiles, from which we isolated *N*_1_ = 50 and *N*_2_ = 500 profiles to mimic a small and a large metabolomic study, respectively. In each case we generated data corresponding to steady state (SS) and non-steady state (NS) solutions. Data were stored in four data matrix 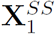, 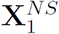 (with dimensions 50 × 83) and 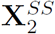 and 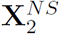 (dimensions 500 × 83. For each of the four data configurations we considered noise levels (SNR) 1%, 5%, 10% and 15%.

### Model representation in matrix form

A network represents relationships among a set of objects, such as the molecular components of a biological system, like genes, proteins or metabolites. The components are represented as as nodes while relationships are represented as links (or edges) between the nodes.

A network can be represented as a square matrix called the adjacency (or also connectivity) matrix. Rows and columns in the matrices represent nodes (in this case, metabolites), whereas non-null entries represent links. In a *weighted* adjacency matrix the entries are real numbers that indicate the strength of the interaction. Values in the *weighted* adjacency can vary, for instance, in the [−1,1] range for correlation, in the [0, +∞) range for mutual information, or in the [0,1] range for probability. A weighted adjacency matrix W can be transformed into a binary adjacency matrix A by imposing a threshold on the (absolute) value on its entries *W_ij_*:

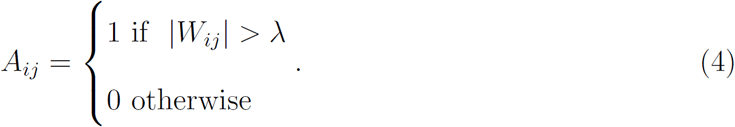

The threshold λ depends on the method considered and on the nature of the adjacency weights; for instance, if correlation is used, λ can be set to 0.6 as proposed in the metabolomics literature;^3^ for mutual information a threshold λ = 0 is usually taken, while for PCLRC, a probability threshold λ = 0.95 has been proposed.^7^

From the network representation of the model (shown in Figure 1), we define three adjacency matrices **A1**, **A2** and **A3** accounting for different degree of association among metabolites, *i.e.* different degree of neighborhood in the AA metabolic pathway.

The three adjacency matrices exemplified in the cartoon shown in Figure 3 and are defined as follows.

**Figure 3.**
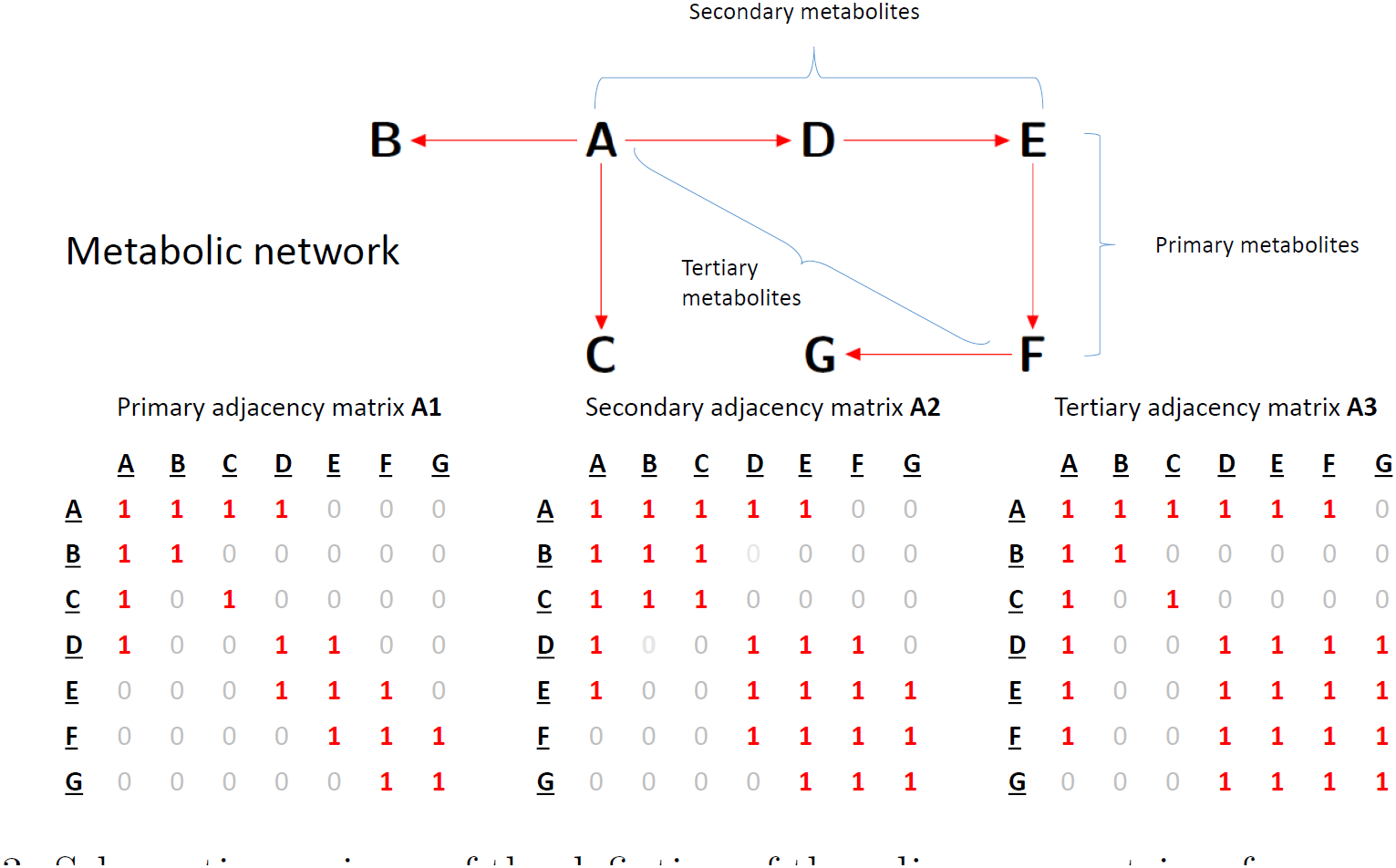
Schematic overivew of the defintion of the adjacency matrices for a metabolic pathway consisting of five metabolites (A, B, C, D, E, F and G) and six reaction (represented by arrows →). Example of primary, secondary and tertiary metabolites are indicated together the corresponding primary (**A1**), secondary (**A2**) and tertiary (**A3**) adjacency (connectivity) matrices. Note that the definition of the matrices takes into account reaction directionality. In this example, **B** and **D** can not be inter-coverted, and as a result they are not connected in any of the adjacency matrices.

i. Primary adjacency matrix **A1**: two metabolites are associated if they connected by a single reaction, i.e they are products or substrates in the same reaction. Examples are arachidonic acid and its direct derivative prostaglandin-H2 (PGH2) and PGH2 and its derivative prostaglandin-H2 (PGE2)

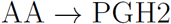
ii. Secondary adjacency matrix **A2**: two metabolites are associated if they are second neighbors in the metabolic map, *i.e.* if they can be inter-converted through two reaction steps via an intermediate metabolite. Examples are arachidonic acid and prostaglandin-H2 (PGE2) connected via PGH2:

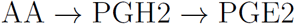 It follows that any two metabolites connected within **A1** are also connected in **A2**.
iii. Tertiary adjacency matrix **A3**: two metabolites are associated if they are third neighbors in the metabolic map, *i.e.* if they can be inter-converted by three reaction steps via two intermediate metabolites. Examples are arachidonic acid and prostaglandin-A2 (PGA2) connected via PGH2 and PGE2:

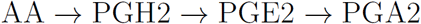 If any two metabolites are connected within **A1** and **A2** they are also connected in **A3**.

### Algorithms for network reconstruction

In our previous work^10^ we have investigated the performance of some algorithm for the reconstruction of metabolite-metabolite associations networks, namely the CLR,^79^ ARACNE,^80^ PCLRC^7^ algorithms together with standard Pearson’s correlations. Here we considered further methods selecting several algorithms from those proposed in the comparative paper by Marbach and co-workers^81^ where over 30 network inference methods were assessed for the reconstruction of gene regulatory networks from high-throughput data. The rationale for the selection was that the methods covered different kinds of approaches and that the algorithms were easily available, easy to use and could be used taking correlation as input metric. It should be noted that, apart from PCLRC, none of the proposed methods was developed with the goal of reconstructing metabolite-metabolite association networks but were developed to study gene regulation. Since in early work we found the resampling approach implemented in PCLRC rather effective, we propose here modifications of existing algorithms by introducing a resampling step. Hereafter, the algorithms considered are briefly explained, mostly following the concise descriptions reported in the supplementary material of Marbach’s paper.^81^ We refer the reader to the original publications for more details.

### CORR - Correlation

This algorithm relies on computing the correlation coefficient among all possible metabolite pairs to infer an association network, although, as previously discussed, correlation may not necessarily translate into association and *viceversa.* The main drawback is the inclusion of chance correlations (especially in the case of small sample size) which may complicate the interpretation of the resulting association networks. Here, we will use both Pearson’s and Spearman’s correlation coefficient.^82,83^ The methods are indicated with CORR-p and CORR-s, respectively.

### PCRLC - Probabilistic Context Likelihood of Relatedness on Correlation

The PCLRC algorithm is based on the CLR algorithm^79^ and has been first introduced to reconstruct metabolite correlation networks.^7^ It combines the CLR approach replacing mutual information with correlation with iterative sampling. In each iteration, 70% of the samples from the full data set are randomly chosen to calculate the correlation matrix. Then, CLR is used to estimate possible associations and reconstruct a network and then dynamical threshold is chosen so that the highest 30% of interactions are kept. The complete procedure (sampling and network generation) is iterated *K* times. The final network is constructed by assigning to every edge a weight defined the frequency of the times that a particular edge was selected in the iterative process. This weight can be viewed as a probabilistic measurement of edge likeliness and can be interpreted as a confidence level on which to accept or reject the correlation between two pairs of metabolites. We set here *K* = 10^5^. The algorithm output a weighted adjacency matrix whose entries are the probability of association for all possible *X,Y* metabolites pairs. We implemented the algorithm with both Pearson’s and Spearman’s correlation: the tow variants are indicated with PCLRC-p and PCLRC-s, respectively.

### MRNET - Multicast Reduction NETwork algorithm

The algorithm works by evaluating the correlation matrix by the Minimum Redundancy Maximum Relevance (MRMR) selection process^84^ for every variable.^15,16^ The logic underlying the MRMR algorithm is that immediate edges are scored higher while distant edges are scored lower. This selection process works by choosing a target metabolite *Y*, and then finding the metabolite, *X_i_*, with the largest correlation with *Y* (maximal relevance). The second selected variable *X_j_* is the one having highest correlation with the target variable *Y* corr(*X_i_*, *Y*) and at the same time, lowest correlation corr(*X_i_*, *X_j_*) with the previously selected variable *X_i_* (minimal redundancy). This set *S* of variables is enlarged by selecting the *X_k_* variables that maximize the MRMR score:

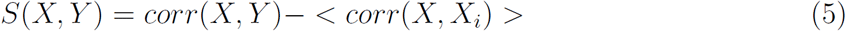

for all *X_i_* already in *S*. The procedure stops when the score becomes negative. The MRMR score represents a trade-off between relevance and redundancy. In practice, for each target variable *Y*, MRNET identifies those variables *X* having maximal pairwise relevance with *Y* and, at the same time, maximal pairwise independence among them.^85^ The algorithm returns a weighted adjacency matrix of the inferred metabolite-metabolite association network: the weight of each pair of metabolites *X* and *Y* is the maximum score between the one computed when *Y* is the target and the one computed when *X* is the target:

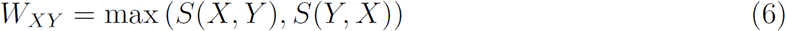

The original MRNET algorithm implemented mutual information instead of correlation. Here we replaced the mutual information matrix with the correlation matrix containing the pairwise correlation among all metabolites in the data set. We implemented the MRNET algorithm with both Pearson’s and Spearman’s correlation, indicated as MRNET-p and MRNET-s, respectively.

### PMRNET - Probabilistic Multicast Reduction NETwork algorithm

The PMRNET algorithm is based on the combination of the MRNETp and MRNETs algorithms and iterative sampling of the data similarly to what done in the PCLRC algorithm. In each iteration, 70% of the samples from the original data set are randomly chosen and MRNETPp or MRNETs is used to calculate the weights matrix *W_XY_* as in Eq. 6). Then dynamical threshold is chosen so that the 30% of the interactions with the highest weights are kept. The complete procedure (sampling and network generation) is iterated *K* = 10^5^ times. The final network is constructed by assigning to every edge a weight defined the frequency of the times that a particular edge was selected in the iterative process. The two versions of this algorithm are indicated as PMRNET-p and PMRNET-s, respectively.

### GENIE3 - GEne Network Inference with Ensemble trees

This algorithm represents prediction of interactions among *p* metabolites as *p* regression problems.^17,86^ Random Forest is then used to predict the concentration of every target metabolite using the concentrations of all the other metabolites. The basic idea of tree-based regression methods is to recursively split the learning sample with binary tests based each on one input metabolite at a time; the tests are optimized to reduce the variance of the output variable (here, the concentration of the target metabolite) in the resulting subsets of samples. At each test metabolite, *K* attributes are selected at random among all candidate attributes before determining the best split. We used 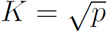 and grow ensembles of 1000 trees.^81^

To each metabolite pair *X,Y* a weight *W_XY_* is assigned that exploits the variable importance measure of the expression of metabolite *X* as derived from the tree-based model learned for metabolite *Y*. For each test metabolite, *X* the total reduction of the variance of the output metabolite due to the split is considered:

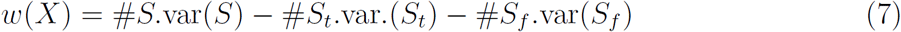

where ♯ is the cardinality of a set of metabolites, *S* is the set of metabolites reaching metabolite *X*, *S^t^* and *S^f^* are the subsets for which test was true and false respectively and *var*() is the variance of the output metabolite in the subsets. The importance measure for all metabolites was calculated as the average of all the scores from all the trees calculated for each metabolites.

### TIGRESS - TRustfull Inference of Gene REgulation using Stability Selection

The TIGRESS algorithm is based on Lasso regression^18^ Similarly to the GENIE3 algorithm every target metabolite is predicted using the concentration of all other metabolites, thus turning the network inference problem into *p* regression problems. Specifically, for each metabolite *Y*, the following regression problem is stated

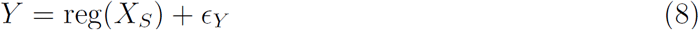

where *Y* is the concentration of the target metabolite, *X_S_* is the subset of candidates metabolites associated with metabolite *Y* and reg() is any (linear or not linear) regression function. A scoring function *g(X)* is defined for each candidate metabolite *X* using Least angle regression (LARS)^87^for feature selection. LARS uses a linear regression model, thus reg() in Equation (8) has the functional form

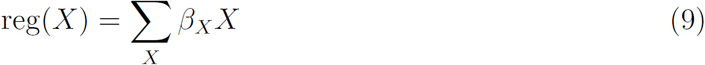

Starting from a constant model where no candidates metabolites are used, metabolites are iteratively added to the model to refine the prediction of the target metabolite *Y*: after *K* = 100 steps of the LARS iteration, a ranked list of *K* metabolites, selected for their ability to predict the target metabolite is obtained. Candidate metabolites are not scored directly using LARS but a stability selection procedure is applied by running the feature selection method many times on randomly perturbed data, and score each metabolite by the number of times it was selected. The input data is perturbed by multiplying metabolite concentrations by a number uniformly sampled in the interval [α, 1] for some 0 < α < 1. Here we used *α* = 0.2 as suggested in the original publication.^88^

This approach has been shown to reduce the sensitivity of LARS and Lasso to correlated features, and improve their ability to select correct features.^88,89^ For each metabolite a final score between 0 and 1 is obtained: a 1 indicates that metabolite *X* is always selected by LARS in the top features to predict the concentration of metabolite *Y*, and 0 indicates it is never selected.^18^

### Metrics for performance assessment

To assess the performance of the different algorithms for the reconstruction of metabolite-metabolite association networks we used the area under the Precision-Recall (PR) curve (also called the Receiver Operating Characteristic curve) indicated with AUC (sometimes also referred as AUROC), comparing the inferred association against the golden standard given by the “true” association network for the AA metabolic pathway as defined by the adjacency matrices **A**_1_, **A**_2_, and **A**_3_ previously defined.

For a binary classifier, the ROC curve is a plot of the recall (*i.e* the sensitivity) against precision (*i.e.* 1 — specificity):

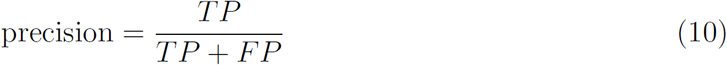

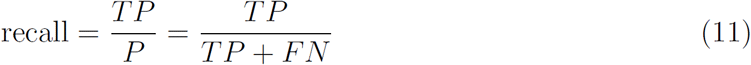

where *P* is the number of positives, *i.e.* the number of interactions in the adjacency matrix representing the model (either **A**_1_, **A**_2_, or **A**_3_); *TP* is number of true positives *i.e.* the number of associations predicted by the algorithm that correspond to interactions in the adjacency matrix representing the model; *FP* the number of false positives: number of predicted associations not found in the adjacency matrix; and *FN* the number of false negatives, the number of associations in the adjacency matrix the algorithm failed to recover.

The methods for network inference considered here produce weighted adjacency matrices, **W**. To compute precision and recall it is then necessary to select a threshold to obtain a binary adjacency matrix (Equation 4). The PR curve is constructed by varying the threshold to take values between the lowest and highest entries in the weighted adjacency matrix. Once the PR curve is plotted, the area under the curve, AUC, is calculated. The limiting value *AUC* = 1 indicates perfect classification, *i.e.* equivalence of **W**with **A1**(or **A2**or **A3**) while *AUC* = 0.5 indicates a random classification.

### Experimental data

Two supporting experimental metabolomics data sets were also considered. The first data set contains 133 quantified metabolites measured on 2139 individuals from the Swedish TwinGene study^90^ using Mass spectrometry. Data were downloaded from the MetaboLights public repository ^91^ (www.ebi.ac.uk/metabolights, accession no. MTBLS93). The second data set contains 29 blood metabolites measured using Nuclear Magnetic Resonance on 864 adult healthy volunteers.^7^ We refer the reader to the original publications for more details on the study design, sample collection, and experimental protocols.

### Software

Calculations were performed using R,^12^ Matlab^11^ and Python.^92^ The R code for PCLRC is available at www.systemsbiology.nl under the software tab. We used the MRNET implementation available in the MINET R package;^93^ and the Bioconductor package GENIE3.^17^ The R code for TIGRESS was downloaded from http://cbio.ensmp.fr/tigress. Simulation and analysis of the AA biochemical network and tits dynamics was performed using COPASI (a COmplex PAthway SImulator).^76^

## RESULTS AND DISCUSSION

### *In silico* generation of metabolites profiles

In this study we wanted to create physiologically plausible metabolite concentration profiles showing correlation patterns similar to those that can be expected to be measured in a standard metabolomic experiment using analytical platforms like Nuclear magnetic resonance (NMR) or Mass spectroscopy (such as LC-MS/MS). Therefore we developed a dynamic model of AA metabolism able to simulate the metabolite concentrations measured in the blood samples of different subjects. The presented model has been parametrized using literature information for kinetic constants and is able to reproduce, at steady state (SS), metabolite concentrations derived from literature.

AA conversion and the formation and degradation of downstream metabolites in the white blood cells and the dynamics of AA derived metabolites in blood (and their sampling) happens at very different scale: hours for blood dynamics and seconds for AA conversion. For this reason, the variation of the downstream products can be considered to be negligible, that is the systems can be considered to be in a steady state. Another reason to investigate the system in steady state is that metabolite-metabolite associations networks are usually build starting from correlations which, in principle, should be calculated starting for samples acquired under the same conditions. If the system(s), in this case, different subjects, are not in the same state, or if they are not in a steady state, the samples (*i.e.* the metabolite concentration profiles) will likely not align, possibly introducing bias in the correlations.^3^

Assuming a complex system like the human body to be in a steady state at the moment of sampling is not always realistic. This assumption may hold true in the case of a few metabolites which are measured under well-defined experimental conditions once the dynamic properties of the system are well characterized, but this is seldom the case in metabolomic studies which usually aim to provide a comprehensive snapshot of the metabolome under given (patho)physiological conditions often without any prior knowledge of the dynamic nature of the systems investigated. For this reason we considered also metabolite profiles sampled in non-steady state (NS), 10 hours after the activation of AA degradation. As expected the individual metabolic profiles acquired when the system is at steady state are more homogenous than those measured in the non-steady state, as indicate by the PCA analysis shown in Figure 4 panel A where the SS profiles are tightly clustered while NS profiles are dispersed in the PCA space.

**Figure 4.**
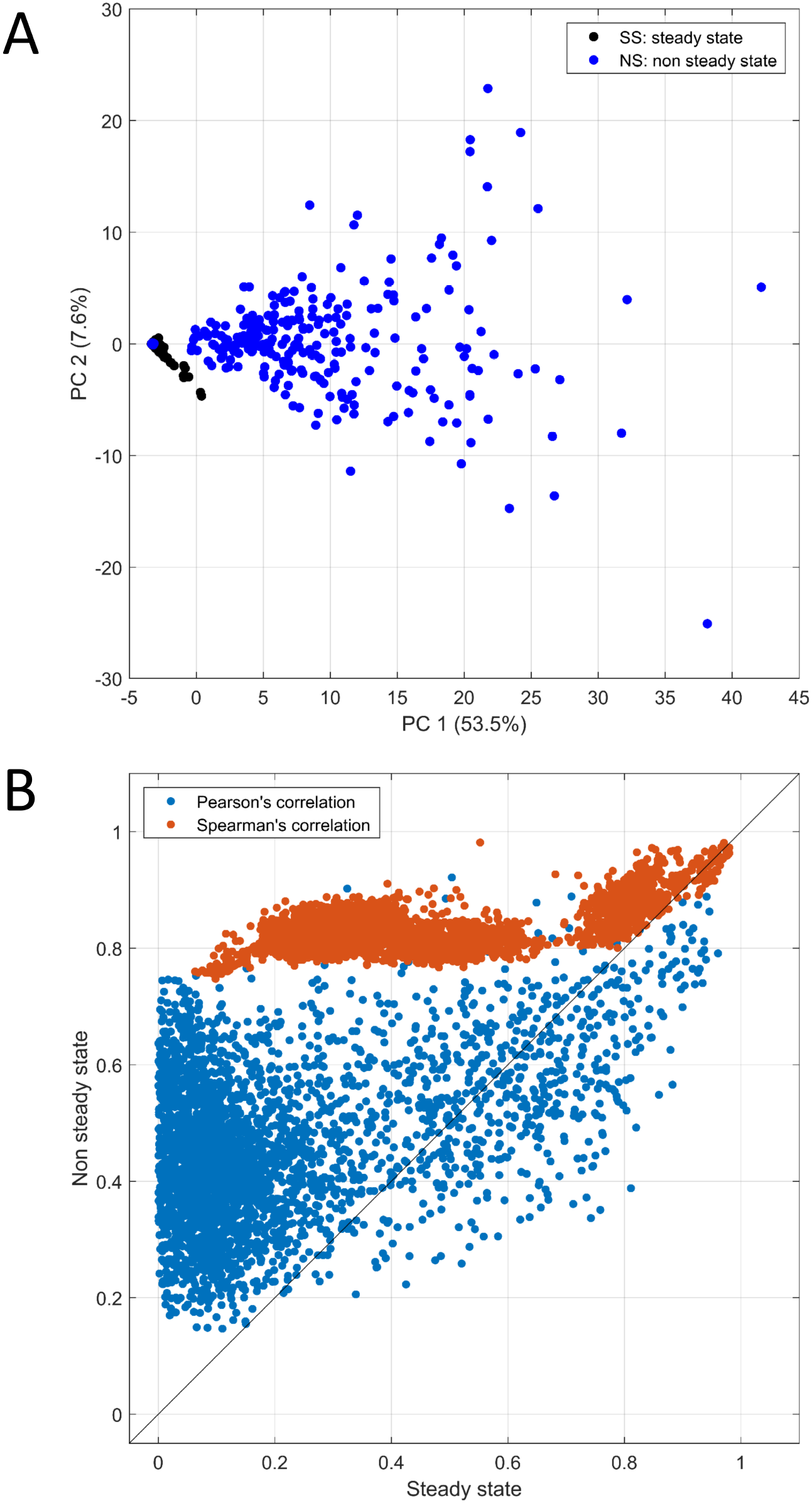
A) Scatter plot of the first two principal components of a PCA model on the data set formed by 500 individual metabolic profiles measure in the steady state (SS) and 500 profiles measured in non-steady state (NS). B) Scatter plot of the (83^2^ — 83)/2 = 3403 pairwise correlations between metabolites for both SS and NS data calculated using both Pearson’s and Spearman’s indexes.

Figure 4 panel B offers a comparison of the metabolite correlation patterns in SS and NS data, calculated using both Pearson’s and Spearman’s indexes. Pearson’s correlations are lower than Spearman’s one, indicating the presence of non-linear relationships among the metabolites: this is what is usually observed in biochemical data because the dynamic governing the relationships among metabolites (see for instance AA model Equation (2)) are highly non-linear.

Correlations in the steady state are mostly below 0.6 (76.8% for Spearman’s correlation and 90.5% for Pearson’s correlations) and this is consistent to what observed in experimental practice and it is supported by chemical considerations: this can happen because the variance in the enzymes that control the reaction involving two metabolites affects them in equal amounts and different directions:^3^ this is what happens to the majority of metabolite pairs in the dynamic model, and is a consequence of the systemic nature of metabolic control.^3^

When correlations are measured on samples acquired at non-steady state we observed a strong increase in the magnitude of correlations: 82.8% of Spearman’s correlation are above 0.8. This may seem counter-intuitive since one can expect that if metabolite concentrations are not aligned the correlation can be biased downwards: however it should be noted that the median variance of metabolite concentrations in the non-steady state is much larger than the median variance in the steady state (3.7 × 10^5^ versus 2.5) and this can bias correlation upwards. Another explanation is that sampling far away from the steady state could results in stronger non-linear relationships, a behavior well described by the Spearman’s index. Another hypothesis is that at the particular sampling point chosen for the non-steady state most part of metabolite pairs are at chemical equilibrium and this usually originates very strong correlations. ^3^

Overall, it seems that data simulated in the steady state shows a correlative structure which is more similar to real metabolomics data: as a term of comparison 93.4% and 91.4% of correlation values (Spearman) observed in two blood metabolomics data sets (MS and NMR, see Material and Methods for more details) are below 0.6.

### Model representation

Three adjacency matrices **A1**, **A2**, and **A3**, were used to represent the original network underlying the AA dynamic. These matrices are schematically represented in Figure 2 and provide standards against which to compare the reconstructed networks. Each of these matrices account for different degree of association (direct and indirect) among the metabolites in the AA metabolic pathway, where association is here defined in terms of distance in the metabolic map.

The classical approach to test algorithms for network reconstruction in metabolomics applications is to generate multivariate data with a given distribution and known covariance-correlation matrix Σ and then reconstruct the correlation patterns which are then compared with the original known correlations. This can be done by directly comparing the actual values of the true and reconstructed correlations (*i.e.* comparing the weighted adjacency matrices) or by comparing the adjacency matrices after that a threshold as been imposed on the correlation values (*i.e.* the patterns of 0 and 1.)

The dynamic metabolic model enables a more exhaustive analysis on the nature of the associations. When metabolite profiles are generated using a dynamic metabolic model, correlations observed in the data do not always reflect the structure of the metabolic network: two metabolites can be direct neighbors in the metabolic network but not correlated; conversely two metabolites can be very distant in the metabolic network but showing high correlation. This a well-known phenomenon and also in the case of this *in silico* simulated data we observe this typical behavior of the dynamic system. For instance, the correlation between PGD2 (prostaglandin D2) and dhkPGD2 (11,15-Dioxo-9S-Hydroxy-5z-Prostenoic Acid) is 0.94 but the two metabolites are separated by two enzymatic reactions and one intermediate metabolite (15kPGD2, 15-dehydro Prostaglandin D2). On the contrary, the correlation among AA and LTA4 (leukotriene A4) is only 0.00015 despite the fact that they are direct neighbors in the metabolic pathway. Another striking example are 6k-PGE1 (6-ketoprostaglandin E1) and 5-oxoETE (5-ketoeicosatetraenoic acid) whose correlation is 0.6 even if they are separated by 8 reaction steps and 7 intermediate metabolites.

From a practical point of view this poses the problem of defining a target to which compare the reconstructed association networks: when are two metabolites to be considered associated in the original metabolic network? Stated otherwise, what is the adjacency matrix corresponding to the AA metabolic pathway shown in Figure 2? This is why we defined three adjacency matrices to represent different degree of association among metabolites in the pathway and this is also the reason we prefer to talk about reconstructed metabolite-metabolite association networks rather than correlation network.

### Performance of network inference algorithms

The performance of the different methods are summarized in Figure 5 for sample size *n* = 500 and in Figure 6 for sample size *n* = 50. Numerical results are presented in the Supporting material S1 (Tables S4-S13).

**Figure 5.**
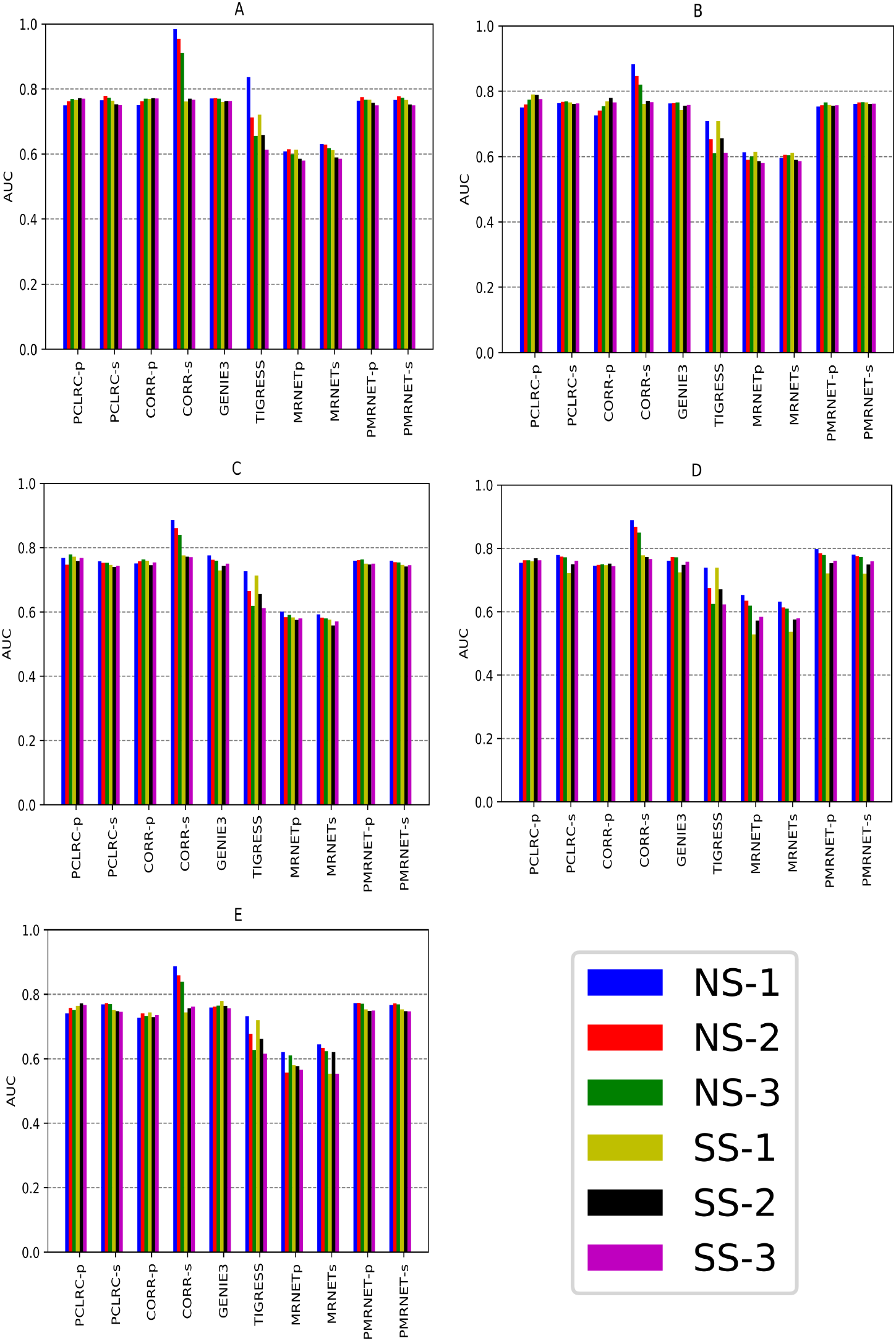
Performance of network reconstruction methods (AUC values) for sample size n = 500 A) no noise data; B) 1% SNR; C) 5% SNR; D) 10% SNR; D) 15% SNR. In color legend, SS and NS indicate that metabolites’ profiles obtained at steady and non-steady state conditions have been used, whereas indices 1,2 and 3 refer to the comparison with the adjacency matrices **A1**, **A2**, and **A3** respectively

**Figure 6.**
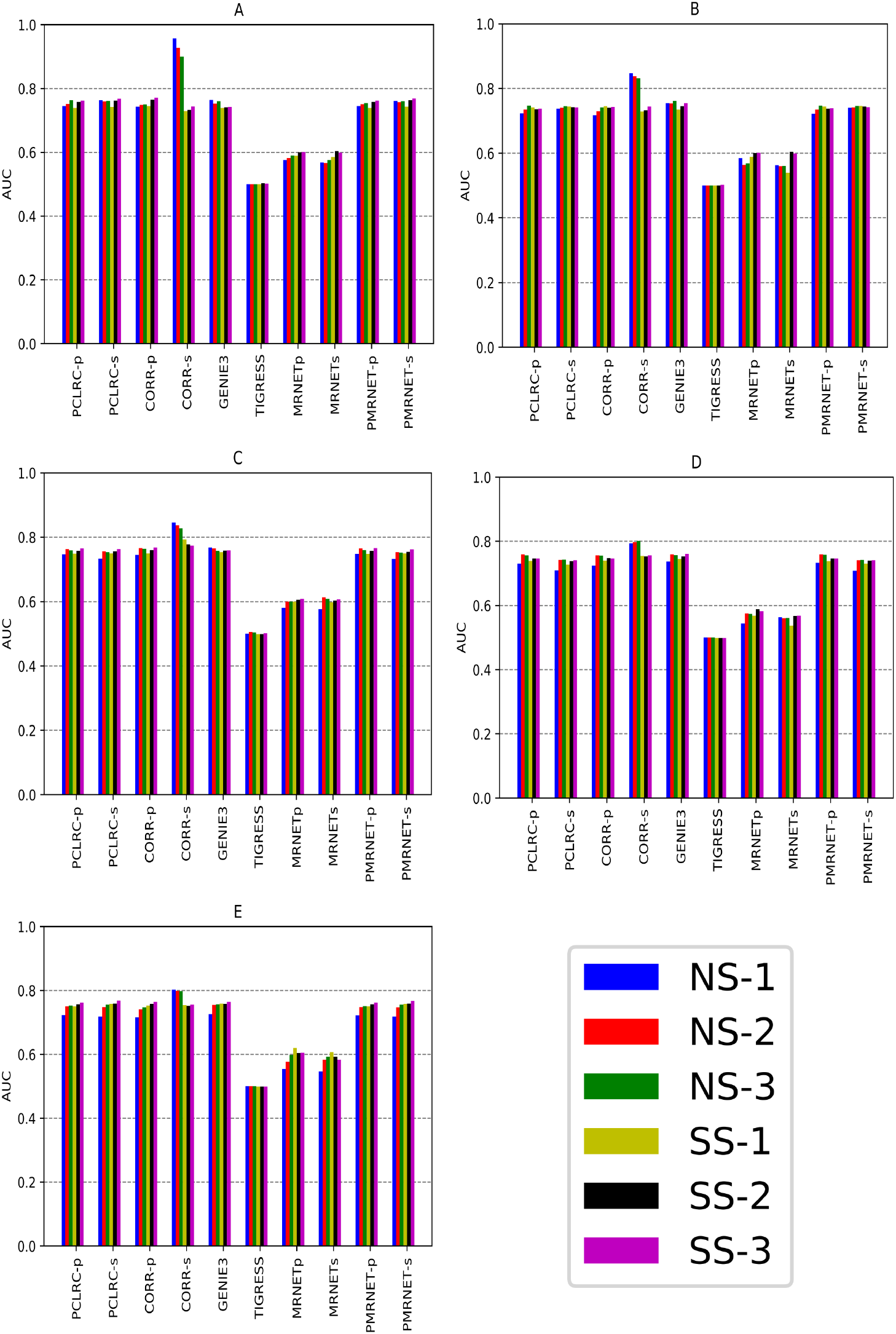
Performance of network reconstruction methods (AUC values) for sample size n = 50 A) no noise data; B) 1% SNR; C) 5% SNR; D) 10% SNR; D) 15% SNR.In color legend, SS and NS indicate that metabolites’ profiles obtained at steady and non-steady state conditions have been used, whereas indices 1,2 and 3 refer to the comparison with the adjacency matrices **A1**, **A2**, and **A3** respectively

The PCLRC algorithm was, overall, the best performer in the comparison, and showed to be robust respect to both sample size and noise: these results are in line with previous results.^10^ The use of simple correlation (either Pearson’s or Spearman’s) is surprisingly good in all situations, which is probably due to the relative larger sample size used, although we note, as expected, a reduction in performance when noise is added to the data.

The performance of the MRNET algorithm was suboptimal in all situations, irrespective of the sample size, data type (SS or NS), noise level or target adjacency used the AUC is constantly below 0.63, indicating that the PR curve is close to the diagonal in the P-R plane, indicating that the solution of the method is insensitive to the particular threshold imposed on the weighted adjacency matrix (*i.e* the λ parameter in Equation (4)) to binarize its entries. This means that the false positive rate is consistently low and the algorithm seems to be able to produce only two types of solutions, namely, with low FPR or the trivial full connected solutions with FPR equal to 1. However, when a resampling approach is implemented (*i.e.* PMRNET-s and PRMNET-p) like in the PCLRC algorithm, the quality of the prediction increase significantly, with increased robustness towards noise.

The TIGRESS algorithm was the most affected by sample size: the AUC values vary between 0.65 and 0.83 for *n* = 500 but dropped to 0.5 for *n* = 50. This result is quite surprising: this algorithm implements a stability selection procedure by randomly perturbing the experimental data by multiplying the metabolite concentration with random numbers. The low performance may be due to the interaction between the small sample size and the perturbation, since a larger sample size is usually needed to compensate for high(er) noise level or to an unfortunate choice of *α*. The algorithm indeed has been found to be very sensitive to this parameter,^18^ however, when perfectly tuned, its performance was found to be equivalent to the GENIE3 algorithm^17,86^ for the reconstruction of gene regulatory networks. Another possible explanation is that the algorithm implements variable selection and it may happen that such procedure is not stable when a small sample size is considered. For the case *n* = 500 the performance of TIGRESS was good although less robust against noise than GENIE3.

The GENIE3 algorithm was the overall top performer in the DREAM3 challenge for gene network inference.^81^ In the present case it also showed a good performance and was minimally affected by changes in sample size. GENIE3 and TIGRESS follow a similar conceptual approach, however in the GENIE3 algorithm regression is performed using Random Forest, that is randomized decision trees, while in TIGRESS linear regression was implemented (see Equation (9)): we have seen that dynamic data exhibits non-linear behavior and Random Forest is particularly well-suited to model non-linear relationships and this may explain why GENIE3 outperformed TIGRESS.

With respect to the use of the primary (**A1**), secondary (**A2**) and tertiary (**A3**) adjacency matrices as target for the network construction, we found the performance of CORR and PLCRC algorithms to increase when considering higher order metabolite-metabolite associations, with PCLRC performing slightly better. The use of PCLRC over the standard correlation approach (CORR) has the advantage that it is not necessary to set a threshold on the correlation value; the weights given to each metabolite-metabolite association do not depend on the magnitude of the correlation itself but on the number of times they appear in the top *Q*% (similarly to what is done in the stability step of the TIGRESS algorithm) and this has the double advantage of *i*) filtering out spurious correlations which may arise from sampling effects and *ii*) maintaining metabolite-metabolite associations regardless from the associated correlation.

This is particularly relevant considering that there isn’t always a correspondence between the correlation among two metabolites and their proximity in the metabolic map. It is of course possible to choose an optimal threshold that simultaneously maximizes precision and recall (see Equations (10) and (11)) but in real life the target adjacency matrix is not known. Other approaches have been suggested, like observing the topology of the network obtained using different thresholds.^94^ Here a transition (from small to large) was observed in the number of connected metabolites at λ = 0.5, a value not far from 0.6, indicated as a lower bound for low/weak correlations in metabolomics.^3^

The performance of the TIGRESS algorithm based on Lasso regression heavily depended on the choice of the target adjacency matrix: when **A1** is used, *i.e.* when only direct neighbors metabolites are considered to be associated, the AUC is 0.836, but it drops to 0.614 when **A3** is considered. This behavior can be explained when considering that this algorithm was developed to infer gene regulatory networks, whose properties are different from metabolite-metabolite association networks. In gene regulatory networks there is asymmetry between the regulators (transcription factors) and their gene targets. This causes genes regulated by the same transcription factors to exhibit similar expression patterns, resulting in a large number of false positives in the reconstructed gene regulatory networks. For these reason algorithms for gene regulatory network reconstruction are often designed to downgrade the importance of indirect associations: this may explain the suboptimal performance of TIGRESS when used to infer indirect associations^1^. However, also GENIE3 has been originally developed to infer gene regulatory networks and, likewise PCLRC, its performance increase when considering higher order metabolite-metabolite associations.

The performance of MRNET is suboptimal in all cases, and it decreases when considering indirect associations, for the same reasons discussed for TIGRESS. However, when a resampling approach is implemented (*i.e.* PMRNET-s and PRMNET-p), the quality of the prediction increases when also considering indirect associations, as observed for PCLRC and TIGRESS.

Regarding the use of Pearson’s or Spearman’s correlations we found in general better perfor-mance when using Spearman’s correlation which is reasonable given the non-linear relationships existing among metabolite concentrations. Overall the performance of all algorithms is reduced in presence of noise: under the additive noise model considered (see Equation (3)) correlations are always attenuated^83, 95^ and this phenomenon is reduced when considering more replicates.

Dependence of the quality of the inferred network on the the use of data acquired in a steady or non-steady state is more complicate to discuss. When a limited sample size is used *n* = 50, we found all methods to perform better on steady state data than on non-steady state and this is reasonable since data is more homogenous as previously observed (see Figure 4.) We also see that for small sample size, the presence of noise has marginally less impact on steady state data than on non-steady state data. When a large sample size is considered (*n* = 500) we do not observe significant differences in the performance of the methods regarding the use of the two types of data. CORR-s performed well in several cases, but it showed the largest decrease in performance depending on sample size and noise level, which may render it sue not optimal for limited sample size.

## CONCLUSIONS

We have explored the ability of several algorithms (and adaptations thereof) to reconstruct metabolite-metabolite association networks using correlation (Pearson’s and Spearman’s) as similarity metrics. We have tested the performance of the algorithms on data generated using a dynamic metabolic model of arachidonic acid metabolism. The model describes 131 reactions interconverting 83 metabolites. The developed model has been parametrized using literature knowledge and is able to reproduce measured concentrations; however it has been greatly simplified and effects such as enzyme or allosteric control have not been explicitly taken into account. This simplification is partly taken into account when considering different degrees of neighborhood in the adjacency matrices representing the model.

While correlation (CORR) is an almost native metric to investigate the association among metabolites in metabolomics studies, all methods but one (PCLRC) were developed to infer gene regulatory networks that have markedly different properties from metabolite-metabolite association networks. We found methods based on resampling like PCLRC (and our modifications of the MRNET algorithm) to perform better together with regression methods based on Random Forest in which resampling is intrinsically implemented.

Except CORR-s in a few cases, none of the methods achieved consistently AUC above 0.8: considering that different adjacency matrices were used to represent the AA metabolic model implementing different levels of neighboring among metabolites and that in general the performance of the methods increased when indirect association are considered we can conclude that not all direct associations (*i.e.* metabolites connected by one single reaction step) are recovered. Rather than from a limitation of the reconstruction algorithms this is an intrinsic property of the systems studied:^3^ due to the systemic nature of metabolic control, the concentration of metabolites connected by one single reaction step may not be correlated and this information cannot be recovered in absence of truly dynamic data or metrics able to capture such behavior.

We are aware that reconstructing and implementing a dynamic metabolic model to generate *in silico* data may not be an easy task and that such approach does not offer the operative flexibility given by standard multivariate approaches, not to mention the lack of statistical characterization of such dynamic data. A possible solution to partially overcome these limitations could be to generate multivariate data with known correlation structure derived from dynamic data and to model data distribution using more advanced data generation techniques.^96,97^

However, it should be remembered that metabolite-metabolite associations are not a property of metabolites and enzymes. Such associations may arise due variation of a single enzyme, or through differential amplification of the variance of a single enzyme but overall metabolite-metabolite association patterns are a property of the whole system. Hence the interest in analyzing biological systems at the network level.

On the basis of this comparative study we recommend the use of inference algorithms based on resampling and bootstrapping when correlations are used as indexes to measure metabolite-metabolite associations. We also advocate for the use of data generated using dynamic models to test the performance of algorithms for network inference since they produce correlation patterns which are more similar to those observed in real metabolomics data.

## Footnotes

1 In a previous study^10^ we observed the same behavior for another algorithm ARACNE^80^ which was not included in the present comparison because of its poor performance.

## SUPPORTING INFORMATION

The following supporting information is available free of charge at ACS website http://pubs.acs.org:

Supporting Material S1. The file contains:

ODE’s for the AA dynamic model

Table S1 : Kinetic values & Reaction rates

Table S2 : All metabolites & their details

Table S3 : All Reactions & their details

Table S4 : AUCs for 500 simulations with no noise

Table S5 : AUCs for 500 simulations with 1% noise

Table S6 : AUCs for 500 simulations with 5% noise

Table S7 : AUCs for 500 simulations with 10% noise

Table S8 : AUCs for 500 simulations with 15% noise

Table S9 : AUCs for 50 simulations with no noise

Table S10 : AUCs for 50 simulations with 1% noise

Table S11 : AUCs for 50 simulations with 5% noise

Table S12 : AUCs for 50 simulations with 10% noise

Table S13 : AUCs for 50 simulations with 15% noise

Supporting Material S2. Arachidonic acid metabolic model in SBML format

## AUTHOR INFORMATION

### Corresponding Author

Edoardo Saccenti; Tel: +31 (0) 3174 82018; Fax: +31 (0) 3174 83829.

### Author Contributions

S. J analyzed data; M.S.-D. and E.S. designed the study; E.S. supervised the work; All authors wrote the manuscript and approved the final manuscript.

### Notes

The authors declare no competing financial interest.

## ACKNOWLEDGMENTS

This work was partly supported by the European Commission funded FP7 project INFECT (contract no. 305340).

